# Discovering paracrine regulators of cell type composition from spatial transcriptomics using SPER

**DOI:** 10.1101/2023.09.27.559863

**Authors:** Tianxiao Zhao, Adam L. Haber

**Author notes:** Correspondence to Adam L. Haber.

## Abstract

A defining characteristic of biological tissue is its cell type composition. Many pathologies and chronic diseases are associated with perturbations from the homeostatic composition, and these transformations can lead to aberrant or deleterious tissue function. Spatial transcriptomics enables the concurrent measurement of gene expression and cell type composition, providing an opportunity to identify transcripts that co-vary with and potentially influence nearby cell composition. However, no method yet exists to systematically identify such intercellular regulatory factors. Here, we develop Spatial Paired Expression Ratio (SPER), a computational approach to evaluate the spatial dependence between transcript abundance and cell type proportions in spatial transcriptomics. We demonstrate the ability of SPER to accurately detect paracrine drivers of cellular abundance using simulated data. Using publicly available spatial transcriptomics data from mouse brain and human lung, we show that genes identified by SPER show statistical enrichment for both extracellular secretion and participation in known receptor-ligand interactions, supporting their potential role as compositional regulators. Taken together, SPER represents a general approach to discover paracrine drivers of cellular compositional changes from spatial transcriptomics.

## INTRODUCTION

Given the specialized character of each cell type, the distribution of types that compose each biological tissue is fundamental to mediating physiological functions. Accordingly, compositional changes, such as influx of immune cells, proliferation of a stem cell pool, or metaplasia of fibroblasts, are tightly regulated by paracrine signaling networks, and these networks are fundamental organizing principles of biological tissues [1]. For example, inflammation can lead to cytokine-mediated changes in cell type composition, such as the canonical ‘weep and sweep’ response to intestinal parasites, involving mucus-producing goblet cell hyperplasia, eosinophilia and basophilia [2]. Many diseases involve this type of change in corresponding tissues: in Alzheimer’s disease for example, neuropathology is associated with aberrant proliferation of microglia and astrocytes [3], which may be mediated by paracrine CX3CL1/CX3CR1 signaling [4]; in asthma, airway remodeling driven by IL-13 leads to increased numbers of smooth muscle and goblet cells accompanied by the abnormal recruitment of immune cells [5]. Triggering factors, like environmental exposures [6] and microenvironmental signals [7], can induce cellular composition changes. To gain insight into the normal physiology of tissue composition and transformations associated with disease, it is essential to understand *which signals regulate the compositions of cell types* in a tissue.

Single-cell RNA-seq (scRNA-seq), combined with computational tools, now provides the transformative ability to simultaneously estimate cell-type composition and measure the expression of factors that regulate it. However, scRNA-seq is unable to determine whether the signals and the receptor cell types are sufficiently near to each other that a putative signaling process is physiologically possible. Spatial transcriptomics (ST) promises to remove this hurdle by simultaneously revealing (1) the expression of paracrine ligands and receptors, (2) the pattern of cell type compositions, and (3) the spatial relationships between the relevant pairs of transcripts and cell populations. ST is an overarching term for a range of methods designed to capture transcriptome-wide gene expression while retaining information about the spatial tissue context [8]. The most widely-used and relatively affordable technologies, including ST Visium [9] and Slide-seq [10], capture spatial information in voxels around 55 microns in diameter, larger than a single cell. Such restriction results in the generation of multi-cell resolution spatial expression data where the absolute number of cells in each voxel is difficult to determine. To address this, recovering the relative proportions of component cell types, can be straightforwardly achieved by available deconvolution algorithms. Probabilistic-based approaches, like RCTD [11], cell2location [12], and SONAR [13], model the expression data with certain distributions under a Bayesian likelihood framework. All these methods produce data which has the constraint that in each spot all proportions sum to 1, and thus their output is compositional. When combined with gene expression data, this type of spatial compositional data can potentially be used to discover paracrine drivers of tissue transformation, but no computational tools yet exist to do so.

In lieu of appropriate methods, several studies have used classical statistical correlations, such as Pearson’s *R* and Spearman’s ρ, to identify such signals in spatial transcriptomics data [14–16]. However, spatial compositional data possesses unique characteristics that requires tailored analytical approaches [17], as treating each individual cell type’s composition as an independent measurement can result in spurious correlations [18]. Additionally, classical metrics face other limitations in this context. For a given cell type *A*, transcriptional signals with spatial dependence on its composition fall into two categories. The first includes *A*-type ‘marker’ genes—those specifically expressed by *A*—which exhibit a trivial spatial relationship with *A*’s composition simply because *A* cells are their primary source. The second, and more intriguing, category includes genes not expressed by *A* cells but whose spatial patterns correlate with *A*’s composition, indicating that they are instead produced by neighboring, non-*A* cells. These genes are potential paracrine regulators influencing *A*’s distribution and abundance.

Machine learning methods have also been applied to spatial transcriptomics to model spatial patterns and cell-cell interactions. For example, MISTy [19] builds multiple components to model the spatial contexts and extracts the spatial relationships between spatial marker genes. SpatialDM [20] uses a bivariate Moran’s statistic to identify the direct spatial association between ligand-receptor pairs, while SpaTalk [21] analyzes cell-level ligand-receptor interactions by testing for enrichment in co-expressed spots. Although these methods are powerful approaches to analyze the spatial distributions of transcriptional signals, most of them, as well as correlation metrics and methods like scHOT [22] and SpatialCorr [23], focus on direct spatial cell-cell interactions and therefore are unable to capture the second category of genes above, as spatial signals in this category may exhibit no direct overlap at all. In addition, none of these methods explicitly model cell type composition, or offer a mechanism to detect drivers of compositional changes.

To overcome this problem, we introduce Spatial Paired Expression Ratio (SPER), a new computational framework to detect paracrine signals regulating the composition of target cell types, even when these signals exhibit non-overlapping spatial dependence. SPER is inspired by the pair correlation function (radial distribution) from physics, which quantifies spatial correlations between two particle types by estimating the likelihood of one particle appearing at a given distance from another [24]. To denoise the spatial compositional signal while appropriately modeling its compositionality, we apply co-kriging, a geostatistical method [25], prior to computing pair correlations to identify the transcripts most strongly co-varying with spatial compositions.

In the following sections, we describe the SPER algorithm and demonstrate, using simulated data, that it substantially outperforms all other applicable metrics: Pearson correlation, Spearman correlation, and Earth Mover’s Distance (EMD). We apply SPER to published ST profiles from adult mouse brain and human lung, and demonstrate that genes identified by SPER are strongly and significantly enriched for transcripts encoding proteins known to be extracellularly secreted, and also known to participate in intercellular receptor-ligand interactions, both consistent with potential roles in paracrine interactions. Moreover, SPER identified novel, biologically significant interactions in the mouse brain data, such as identifying the neuropeptide precerebellin (*Cbln1*) as a novel factor associated with abundance of GABA-ergic Meis2^+^ neuron, which express its cognate receptor *Nrxn2*, as well as the Wnt ligand R-spondin 3 (*Rspo3*) associated with the same Meis2^+^ neuron subset, which also express its receptor *Lgr5*. In human lung data, SPER detects the fibroblast-derived signal fibrillin-1 (*FBN1)* associated with alveolar type II (AT2) cells, which express the integrin αvβ6 (*ITGB6*), one of the known receptors for FBN1. These novel putative paracrine interactions detected by SPER were not identified using other available computational methods. Taken together, our findings establish SPER as a general and effective approach for identifying putative regulators of cell type composition from spatial transcriptomics data.

## RESULTS

### SPER scores paracrine signals by evaluating spatial dependence between genes and cell types

Spatial Paired Expression Ratio (SPER) is a computational framework designed to uncover paracrine signals that regulate cell type compositions within spatial transcriptomics datasets. Unlike classical correlation-based approaches that are prone to detecting trivial spatial associations, SPER explicitly models the spatial interplay between genes, potentially encoding paracrine signal molecules, and cell types within a certain distance. Studies suggest that paracrine signals typically function within a maximal distance of 25 cell diameters, approximately 250-300 µm [26]. To detect these interactions, SPER employs the pair correlation function that can describe how spatial correlation changes as a function of distance. For every gene-cell type pair, SPER evaluates how the dependence of the gene’s expression level on the target cell type’s abundance varies with distance to the target cell type and identifies those pairs showing strong a signal of paracrine spatial dependence.

The method integrates three key components: spatial gene expression, spatially resolved cell-type compositions (derived from deconvolution algorithms), and spot-to-spot spatial distances (**Fig. 1**). As the first step, SPER applies geostatistical techniques, specifically co-kriging, to de-noise compositional data while accounting for inherent spatial dependencies. For each gene-cell type pair, it then computes the gene’s expression ratio to its global average expression level, defined as ‘paired expression ratio’, across different distance lags. The outcome, as a paired expression ratio spatial density distribution, will be convolved with a spatial weight into one score to prioritize biologically relevant interaction scales. After adjusted with the expression prevalence in the target cell type, a SPER score matrix can be eventually calculated based on every pair of gene and cell type. This workflow enables SPER to differentiate between direct, overlapping spatial associations and more subtle, non-overlapping dependencies indicative of paracrine regulation.

**Figure 1.**
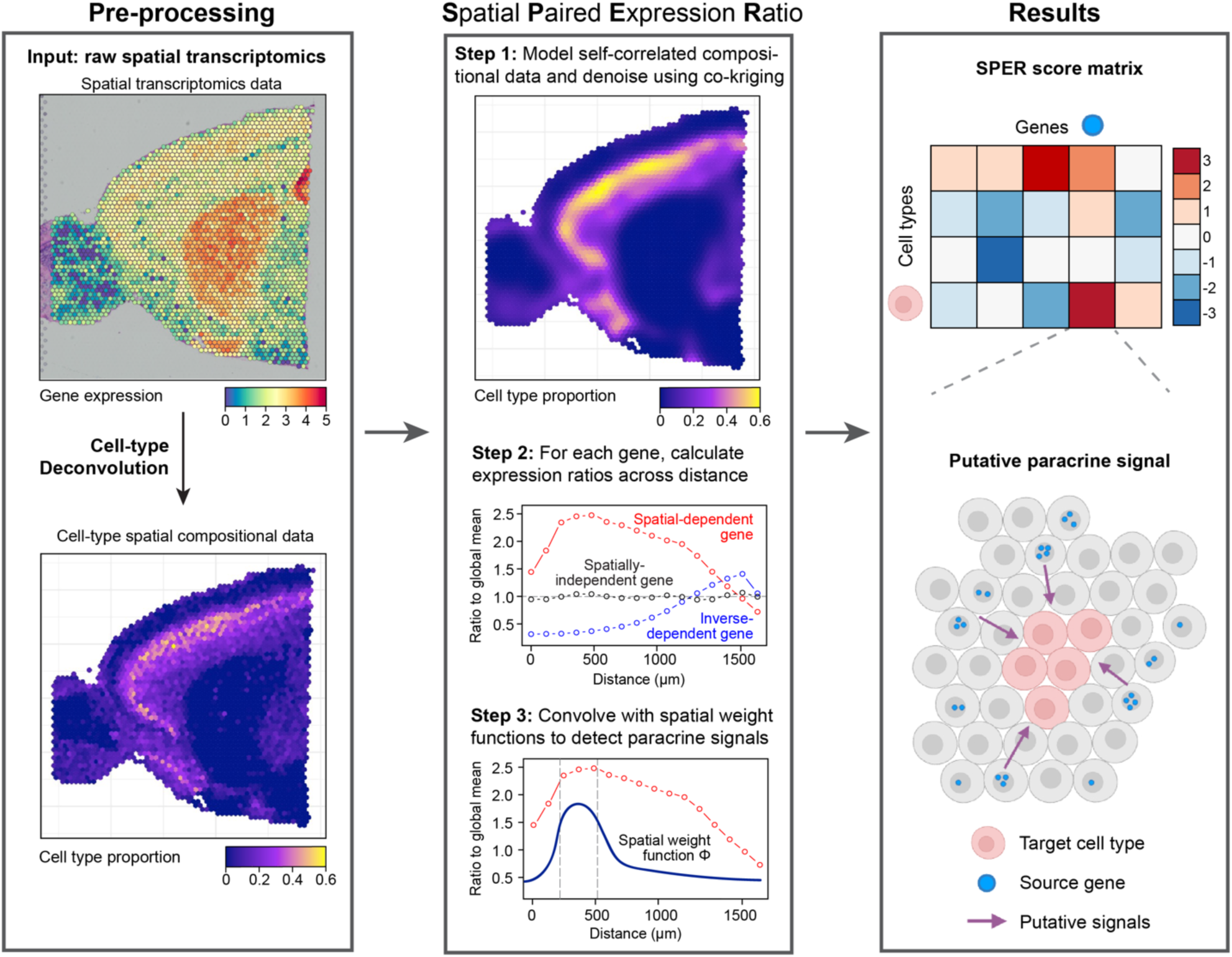
Schematic of spatial paired expression ratio (SPER) algorithm. The workflow of SPER: (Left) **Pre-processing**: SPER requires three different inputs: spatial transcriptomics data, spatial cell-type compositional data, and the spot-spot distance matrix. Any cell-type deconvolution method can be used to generate the spatial compositional data. (Center) **Core algorithms**: SPER models the spatial dependence between genes and their potential target cell types by measuring the expression changes over the interaction distance of a cell type-gene pair. As the first step, we apply co-kriging algorithms to model and de-noise the spatial compositional data. Next, we evaluate the spatial association by calculating a gene’s ratios to its mean expression over the distance towards the target cell type. These ratios are defined as the paired expression ratios, and a spatial weight function on these ratios is applied at the last step to emphasize paracrine signal patterns. (Right) **Results**: the algorithm generates a SPER score matrix as the final output, where higher scores indicate a stronger confidence of this cell type-gene pair as a putative paracrine signal, with a schematic example transcript cell-type pair shown (bottom right).

While no method yet exists to mine ST data for paracrine signals regulating cell type composition, the major analytical field to derive inferential tools for spatial compositional data has been geostatistics. This branch of statistics focuses on modeling multivariate data that show codependency related to spatial proximity, originally developed in predicting the co-distribution of metal ores for mining applications [27]. Co-kriging is a geostatistical interpolation approach that models’ compositionality as well as spatial dependance between components of a mixture. It uses kernel functions to describe the pattern of variograms between each pair of components as a function of geographic lag distance [18]. SPER leverages these desirable properties of co-kriging to de-noise the spatial compositional data, modeling variograms between cell type proportions, and adjusting for spatial autocorrelation. Then, using the distance information obtained from the physical coordinates of all ST spots, calculates the paired expression ratio as a series of matrices, which includes all pairs of cell types and genes; finally, SPER convolves the ratios at each distance with a weight function to integrate over length scales of interest and obtain a scalar score representing the spatial relationship between the given cell type and the gene (**Fig. 1** and **Methods**).

### SPER detects simulated paracrine drivers of cell type composition

To evaluate how well SPER detects paracrine regulators of cell composition, we applied it to three different simulated ST datasets (**Fig. 2**) and compared it with traditional methods of detecting spatial co-variation, specifically: Pearson correlation, Spearman correlation, and Earth Mover’s Distance (EMD) [28]. To set up these synthetic benchmark datasets, we simulated 80-by-80 grids where each tile represents a spot in the physical space. Next, different cell types and gene expression patterns were assigned to tiles in the following manner: cell type specific ‘marker genes’ for each cell type, only expressed in the corresponding cells, were defined across the space; the ‘paracrine signal genes’, which we are interested in, can be mathematically described as genes having a similar but not directly corresponding spatial distribution as their target cell type. Such genes can be expressed either by a sole source cell type or be jointly expressed by multiple cell types. Therefore, we setup three different scenarios to simulate the paracrine signals: simplified single-source, realistic single-source, and realistic multi-source paracrine signal patterns (**Fig. 2, left**), using radially dependent association for a simplistic and Markov random fields for more realistic simulations (**Methods**). With these simulated ST data, cell-type deconvolution methods were then applied to generate spatial cell type compositional data. SPER, as well as the other canonical statistical metrics, were then applied to detect potential paracrine signals affecting target cell composition.

**Figure 2.**
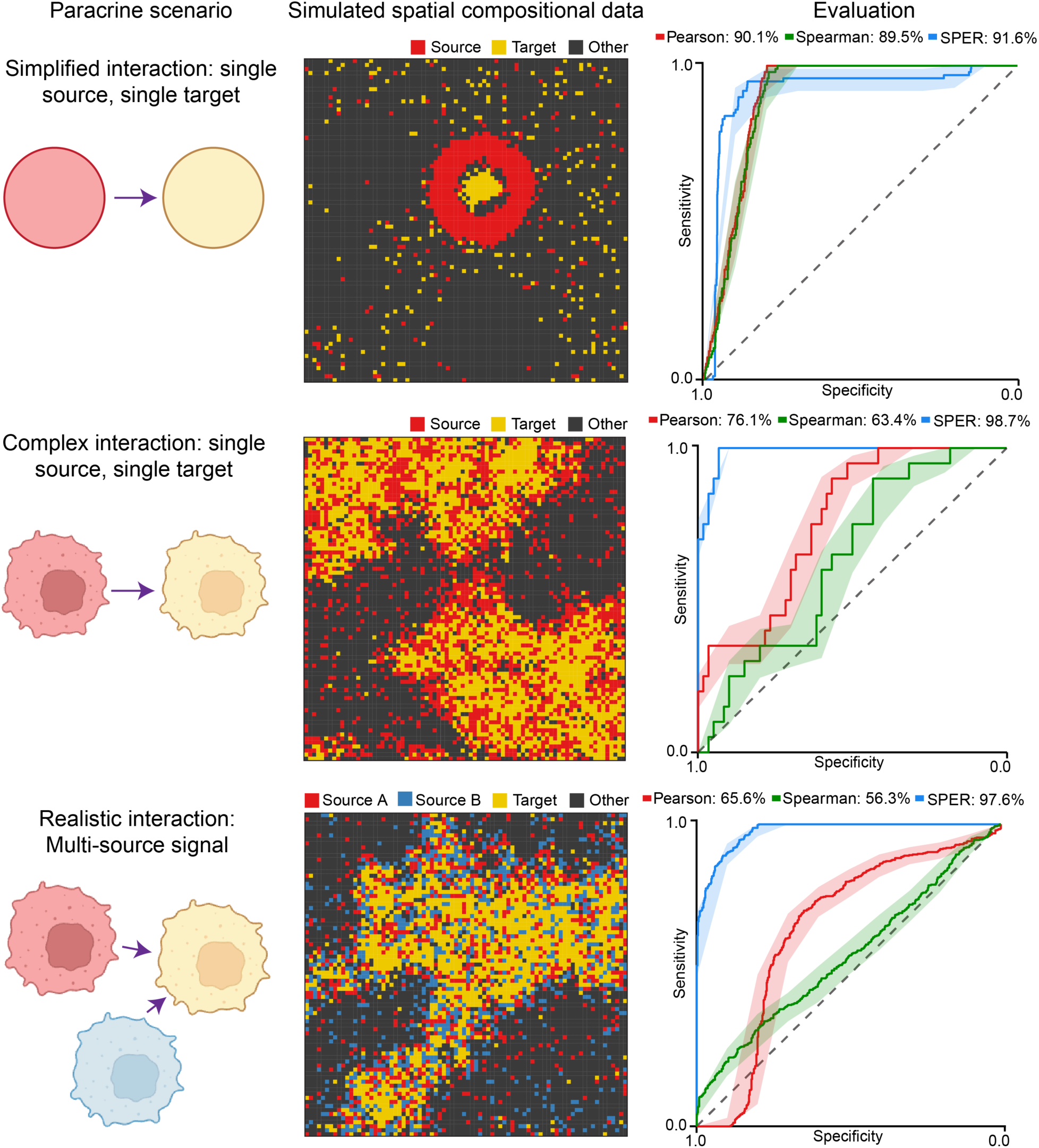
Evaluation of SPER on different simulated spatial transcriptomics data. **Left column**. Schematic of the three paracrine signal patterns - simplified single-source, realistic single-source, realistic multi-source - simulated in this study. **Middle column**. Visualizations of representative simulated spatial compositional data for each pattern type (rows) in an 80×80 grid, with each tile colored by its major cell type (color legend). In the realistic scenarios (bottom two rows), Markov random fields were used to simulate the spatial dependence between the sources and target. **Right column**. Area under ROC curve (AUROC) values are calculated to evaluate how well each metric distinguishes simulated paracrine signals from other spatially unrelated genes.

In the simplified single-source simulation, paracrine signal genes are expressed by a single source cell type located at a fixed average radial distance of 10 units from the target cell type. This setup ensures that the spatial distributions of signal genes spatially depend on, *but do not substantially overlap with*, the distributions of the corresponding cell types (**Fig. 2, top row**). We reasoned that this lack of overlap would be characteristic of a paracrine signaling interaction where a neighboring cell population provides a ligand to a target cell population. To enhance the biological fidelity of the synthetic data, we developed a more realistic single-source simulation where the spatial distribution of cell types is modeled using a Markov random field [29]. In this scenario, target cell abundance is spatially associated with, but again remain distinct from, the abundance of their simulated paracrine ligand in a more biologically realistic arrangement (**Fig. 2, middle row**). Furthermore, to account for signaling pathways involving multiple sources, we simulated a multi-source scenario in which target cells are surrounded by two source cell types expressing the signal genes (**Fig. 2, bottom row**).

To compare the paracrine pattern to other genes, we also simulated a control set of genes that are uniformly distributed in the grid and therefore have no spatial association with any cell type. We considered the marker genes expressed by target cell types as part of control group. Thus, we can evaluate a method’s performance on detecting paracrine gene signals by how well it can distinguish them from the control set. Four different metrics, SPER, Pearson and Spearman correlation, and Earth Mover’s Distance (EMD), were applied in the simulations (**Fig. 2, right**). While all other metrics completed in minutes, EMD, a method with extremely high computational complexity, failed to complete within two hours. Therefore, we decide to exclude this metric from the list as it is therefore impractical to apply on real data.

We repeated each simulation 10 times to generate robust results. Given the output scores from each method, we calculated the ROC curve of discriminating the paracrine signal genes from other genes. While all three methods exhibited a good detection accuracy in the simplified simulation, SPER consistently showed the best performance, accurately detecting signals in all three scenarios with AUROC scores from 0.92 to 0.99. Pearson and Spearman correlations, which did well in the simplified scenario, failed to capture the paracrine signals in two more realistic simulations with the AUROC scores from 0.56 to 0.76. Taken together, our simulation studies demonstrate the ability of SPER to detect paracrine signals that might be missed by other traditional metrics, especially in a more complex and realistic settings.

### SPER outperforms classical metrics in detecting potential paracrine signals in real ST data

To evaluate its performance on real data we next applied SPER and the other metrics to ST profiles of mouse brain generated using 10X Visium [9]. After preprocessing, 18270 unique genes were detected across 2695 spots sampled from a sagittal anterior mouse brain section. Based on benchmarking studies evaluating performance on this task [30–32], we selected RCTD [11], which consistently ranks among the top-performing methods, in this study to estimate the cell type composition with matched scRNA-seq as the reference [33]. (**Fig. S1A**). SPER’s co-kriging models showed good fits to the observed variograms of top ten cell types (**Fig. S2**), and visualizing the compositional data after co-kriging demonstrated its ability to remove noise (**Fig. S1B**). For each of the ten neuronal cell types in the dataset, we used a published scRNA-seq reference dataset to define cell-type-specific marker genes that were strongly upregulated in one particular cell type (at least two-fold higher, FDR<0.001, calculated using MAST [34]). Given the cell type source of these transcripts, we expect their expression distribution in the ST data to be spatially strongly coupled to the abundance of that cell type. Using Pearson and Spearman correlations, we confirmed that these marker genes showed significant spatial dependency to their corresponding cell types (**Fig. S3**).

However, our goal is to identify paracrine signals, we hope to detect those genes that are not expressed (or only lowly expressed) in the target cell type but are instead expressed by neighboring source cells. We defined ‘putative paracrine transcripts’ as those whose spatial dependency was strongly associated with a target cell type but had only low expression in that target cell type. To maximize detection of these transcripts associated with—but not expressed by—a cell type, the SPER score also considered the gene expression prevalence matrix from the scRNA-seq expression data (**Methods**). To visualize the extent to which putative paracrine transcripts were enriched in each metrics top ranked genes, we plotted the metric scores against the proportion of cells expressing each gene in the reference scRNA-seq data [33] for each cell type-gene pair (**Fig. 3A-B, S5**). In L4 (Layer 4) neurons for example (**Fig. 3A-B**), all genes above the 95^th^ percentile of Pearson correlation also showed high expression prevalence in L4 neurons; and so, it detected just 3 putative paracrine signals whose expression prevalence less than 0.5 at the 95^th^ percentile threshold (**Fig. 3A**), indicating that virtually all transcripts associated with L4 neurons are expressed by them, and many of them are trivially L4 marker genes. On the other hand, in the case of SPER, there are no marker genes within the top 95^th^ percentile of association scores, and the majority meet our baseline criterion to be putative paracrine transcripts, which we define as genes whose expression prevalence is less than 20% in the target cell type (**Fig. 3B**). This trend was consistent across all cell types after varying the metric percentile thresholds from 75^th^ to 95^th^, where SPER detected markedly more putative paracrine genes meeting this criterion (**Fig. 3C**), supporting SPER’s capability to detect putative paracrine regulators of cell type composition.

**Figure 3.**
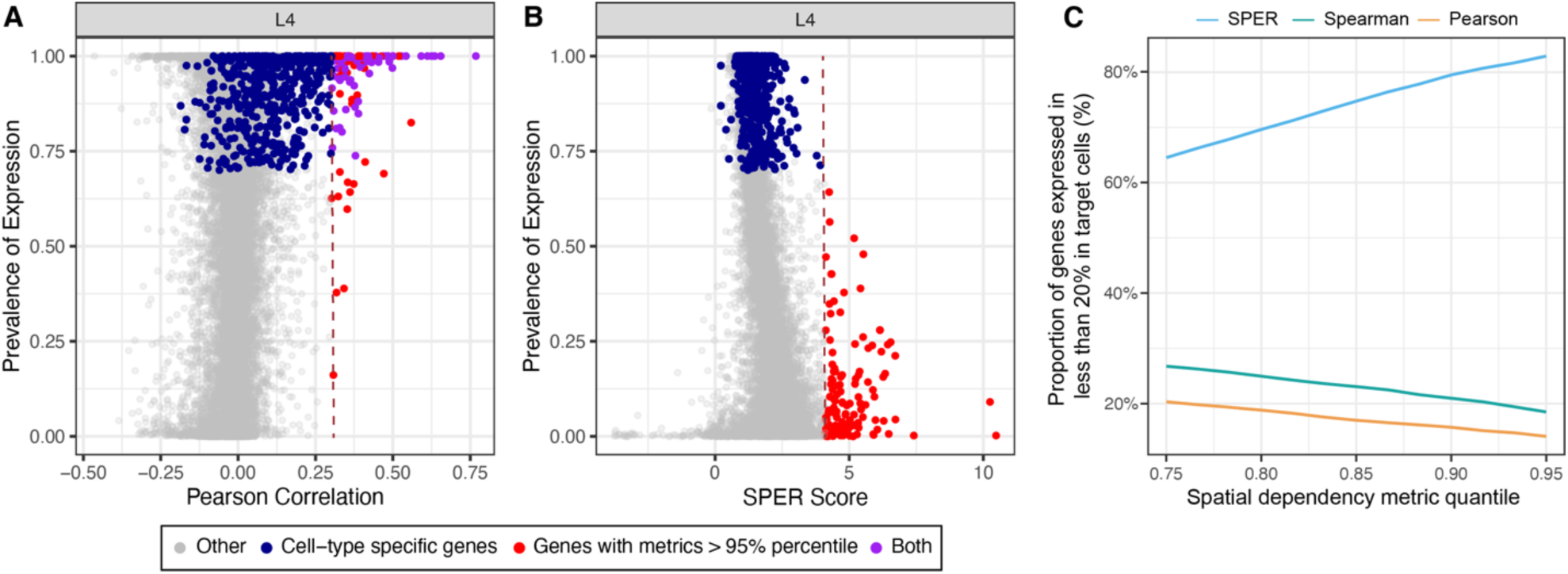
SPER detects non-trivial spatial relationships. **A-B**. Scatter plots show the distribution of Pearson correlations (A) and SPER scores (B) with the proportion of layer 4 (L4) cortical neurons in which each gene (dot) is detected in scRNA-seq profiles. The dark blue dots mark the marker genes for each cell type (**Methods**). Red dots stand for the genes whose metric scores are above the 95% percentile. Purple dots represent genes that meets both conditions. (C) Proportion of top-ranked (above the given percentile, x axis) genes that are not trivially expressed by the target cell type (defined as expression prevalence less than 20% in the target cell type) for each metric (color legend, top).

### Transcripts with high SPER scores are enriched for extracellular proteins and paracrine ligands

If the genes spatially related to cell type composition are indeed participating in paracrine interactions, then we reasoned that they must encode proteins capable of performing these functions in facilitating intercellular communication. We therefore tested whether the putative paracrine transcripts identified by SPER and other methods were statistically significantly enriched for (a) genes known to be secreted into the extracellular space and (b) genes known to act as paracrine ligands (**Fig. 4**). As before, we set thresholds on the metrics, requiring them to have a SPER or other metric score larger than the 95^th^ percentile of all genes. As expected, transcripts identified by SPER were significantly enriched (*p*<0.001) for both extracellular genes (**Fig. 4A**) and paracrine ligands (**Fig. 4B**) for all ten cell types in the mouse brain dataset. The other two metrics were significantly enriched (*p*<0.05) only in three and four cell types for Pearson and Spearman correlation respectively for extracellular genes (**Fig. 4A**), and in four and three respectively for known paracrine ligand genes (**Fig. 4B**). With few exceptions, SPER identified more of both types of transcripts compared to Pearson and Spearman correlations (**Fig. 4**). Lastly, we performed a sensitivity analysis to confirm the utility of co-kriging in denoising our compositional data (**Methods**). We found that while transcripts highly ranked by SPER without co-kriging still showed enrichment, the addition of co-kriging improved the enrichment for both extracellular location and known paracrine ligand activity for the majority of cell types, demonstrating its utility in improving SPER’s sensitivity (**Fig. S4**).

**Figure 4.**
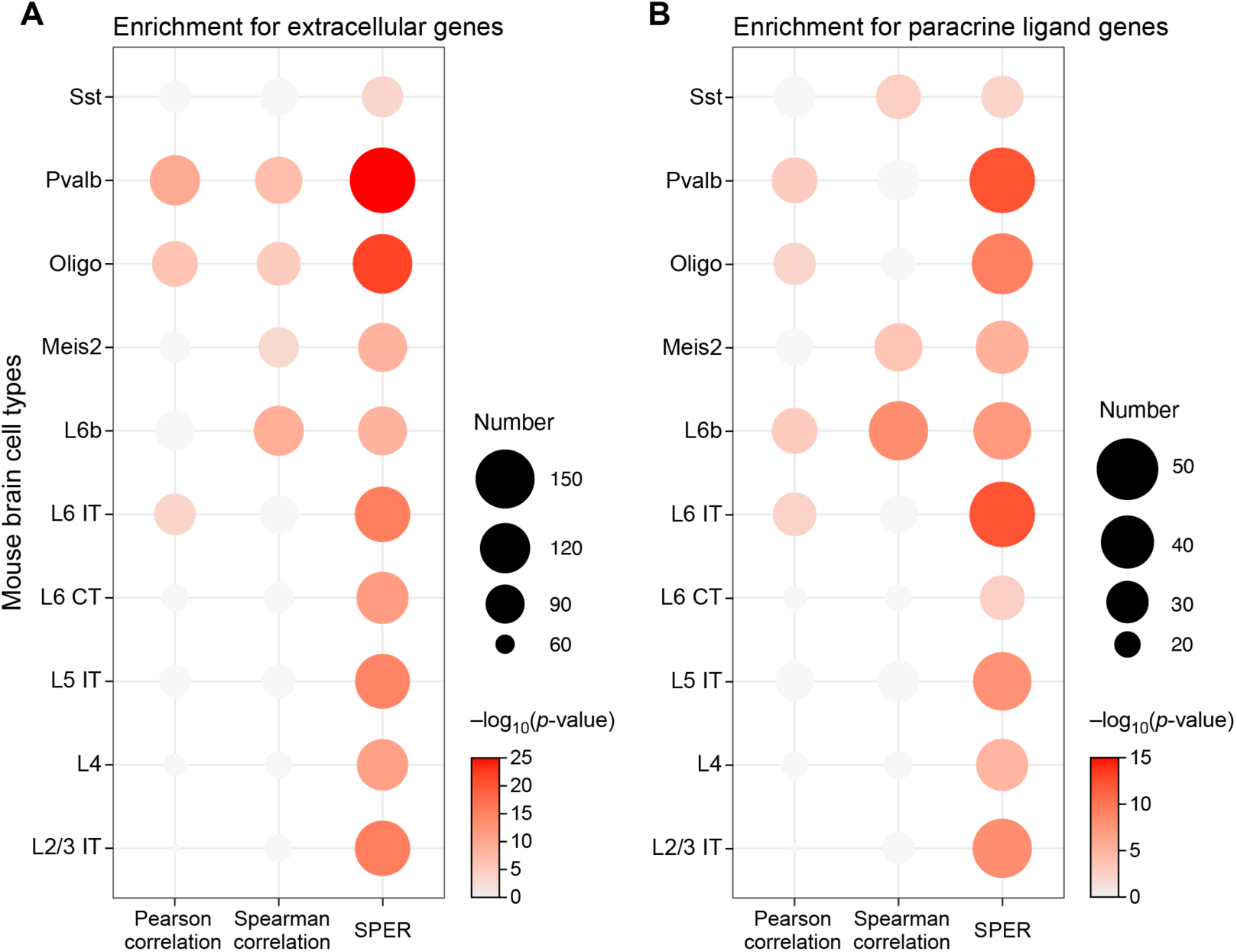
SPER’s top-ranked genes are enriched for both extracellular genes and known paracrine ligands. Dot plots visualize hypergeometric enrichment analysis of extracellular (A) and known paracrine ligand signals (B) within candidate gene sets. Extracellular and ligand genes annotations were collected from COMPARTMENTS, FANTOM5, and CellPhoneDB databases respectively (see **Methods**). The candidate sets include genes whose scores for each spatial dependency metric (x-axis) are above than the 95% percentile of all genes. The size of dots represents the number of extracellular genes or ligand genes in the given gene set. The color represents the significance the enrichment in the set (hypergeometric test), non-significant (*p*>0.05) overlaps are shown as light grey.

Together, these results demonstrate that candidate paracrine transcripts identified by SPER have three important characteristics: (1) they are predominantly expressed by a neighboring cell type, not the target cell, but they are nonetheless spatially related to the target cell’s composition, and they are also statistically enriched for (2) genes known to be secreted into the extracellular space, and (3) genes known to participate as ligands in paracrine interactions.

### SPER detects single- and multi-source putative paracrine signals in mouse brain

Using SPER, we have detected a set of putative ligand genes that have high spatial association with the composition of a putative target cell type. Across all cell types there are 5569 pairs of cell types and putative paracrine regulators above the 95^th^ percentile of SPER scores. Of these, 250 also showed substantial expression of the cognate receptor in the target cell types: a total of 506 ligand-receptor pairs, each potentially active in regulating the composition of the brain tissue in the sample. For example, *Cbln1* (precerebellin) and Meis2^+^ neurons (a cell type defined by expression of the homeobox transcription factor *Meis2* [33, 35]) showed a strong spatial association, but also non-overlapping spatial distribution, indicating possible paracrine compositional regulation (**Fig. 5B**). Notably, it was not identified by Pearson correlation, ranking 349^th^ (*R*=0.32). On the other hand, using SPER we found that the *Cbln1*-Meis2^+^ neurons pair had the 4^th^ highest score among all Meis2^+^ neuron-related signals where the appropriate cognate receptor was also expressed. Consistently, the scRNA-seq data showed that *Cbln1* is only lowly expressed in Meis2^+^ neurons, but is indeed highly expressed by the neighboring L6 IT neurons (**Fig. 5A**). In this case, reference scRNA-seq data showed that the paracrine signal is expressed only by this single source cell type. We examined *Nrxn2*, a receptor for *Cbln1* (**Fig. 5C**), and found as expected that it was expressed on Meis2^+^ neurons, supporting the possibility of Cbln1 acting on them. In the cerebellum, *Cbln1* is known to play an important role in the Purkinje cell synapse formation [36] as well as the growth and guidance of axon in multiple neural regions within mouse embryos [37], but to our knowledge, its role as a paracrine signal in the motor cortex has not yet been described.

**Figure 5.**
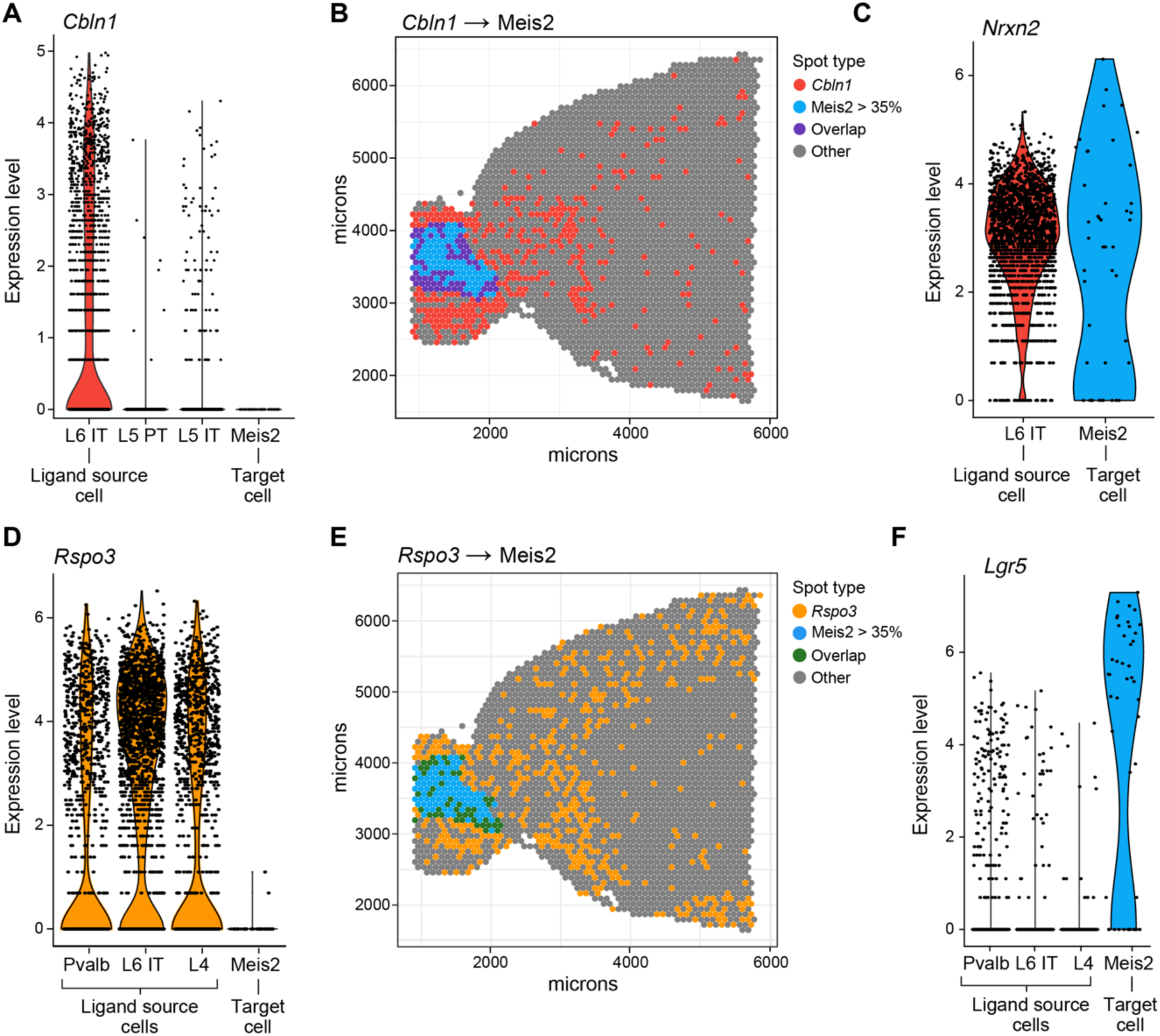
SPER detects single-source (*Cbln1*) and multi-source (*Rspo3*) paracrine signals in mouse brain. (**A, D**) Violin plots show expression levels (y-axis) of *Cbln1* and *Rspo3* in relevant mouse brain cell types (x-axis) from reference scRNA-seq data. L6 IT, L5 PT, L5 IT, L4, and Pvalb neurons are the neighboring cell types of Meis2^+^ neurons. (**B, E**) ST plots visualize the spatial dependence between transcript-target cell type pairs *Cbln1*/Meis2^+^ neurons (B) and *Rspo3*/Meis2^+^ neurons (E). The color of spots (color legend) shows whether *Cbln1* (B) or *Rspo3* (E) is detected, a proportion of Meis2 cell larger than 35%, both (overlap), or otherwise. (**C, F**) The expression levels of the relevant cognate receptor *Nrxn2* (C) and *Lgr5* (F) in the reference scRNA-seq data.

Paracrine signals can of course also be derived from multiple cell types. For example, SPER identified *Rspo3,* which encodes the Wnt ligand R-Spondin 3. *Rspo3* is produced by several source cells: Pvalb^+^, L6 IT, and L4 neurons (**Fig. 5D**), and shows a non-overlapping spatial association with the identified target cells, again Meis2^+^ neurons (**Fig. 5E**). While *Rspo3* is identified as a significant putative signal by SPER, it cannot be found as a paracrine signal using any correlation metrics (**Supplementary Table 1**). We tested for expression of the cognate receptor for *Rspo3* in Meis2 cells and found that *Lgr5* (Leucine-rich repeat-containing G-protein coupled receptor 5) was strongly and selectively expressed in the target cell type, consistent with an active paracrine interaction (**Fig. 5F**). This suggests that *Rspo3*, released by Pvalb, L6 IT and L4 cells, may serve as a paracrine regulator to Meis2 cells, activating the *Wnt* signaling pathways, which could potentially shape the spatial compositions of Meis2^+^ neurons. While this interaction is well-studied in the intestine [38], to our knowledge its role in the mouse motor cortex is not yet known. The link between *Rspo3* and Meis2 cells, therefore, provides a representative example of multi-source paracrine compositional regulators that can be discovered using SPER.

### SPER enables discovery of paracrine signals from established human lung spatial dataset

To validate SPER’s sensitivity to detect such paracrine signals, we also applied it to a second published ST dataset of the human lung [39]. The processed Visium spatial transcriptomics data was deconvoluted by cell2location [12], based on the paired reference scRNA-seq data with a total of 1147 genes in approximately 200 thousand cells. The deconvolved ST data (**Fig. 6**) contains 3234 spots and the relative compositions of 80 annotated cell types. We applied SPER (Methods) to identify potential paracrine regulators of human lung cell type abundances (**Supplementary Table 2**). In addition to identifying many known paracrine interactions—such as club cell-derived *SCGB3A2* and alveolar macrophage abundance, via the *MARCO* receptor [40] or basal epithelial cell-derived *CXCL14* and B cell abundance, via the receptor *CXCR4* [41]—SPER identified a novel fibroblast-derived signal: fibrillin-1 (*FBN1*), an extracellular matrix (ECM) protein [42], was spatially associated with the composition of alveolar type II (AT2) cells (**Fig. 6A-B**). The association for FBN1 and AT2 cells was very strong, ranking in the 98^th^ percentile of SPER scores (20^th^ of 1147 tested genes), but both Pearson and Spearman correlation scores failed to capture this signal with *R*=0.056 and *ρ*=0.061, outside the top 200 ranked transcripts for both metrics. Consistent with a plausible paracrine signaling role of fibroblast-derived FBN1, integrin αvβ6 (*ITGB6*), an epithelial-exclusive integrin that acts as a receptor for FBN1, is strongly expressed on AT2 cells (**Fig. 6C**). In lung organoid models, diverse ECM proteins including FBN1 are known to be critical for differentiation and proliferation of AT2 cells [43], but to our knowledge, the direct link between fibroblast derived *FBN1* and *ITGB6* on AT2 cells has not been previously identified as active *in vivo*. Mutations in *FBN1* result in increased availability of TGFβ and are linked to the congenital disease Marfan’s syndrome, the effects of which are predominantly cardiovascular, skeletal, and ocular, but also induce a respiratory phenotype [44] including lung lesions and spontaneous pneumothorax [42].

**Figure 6.**
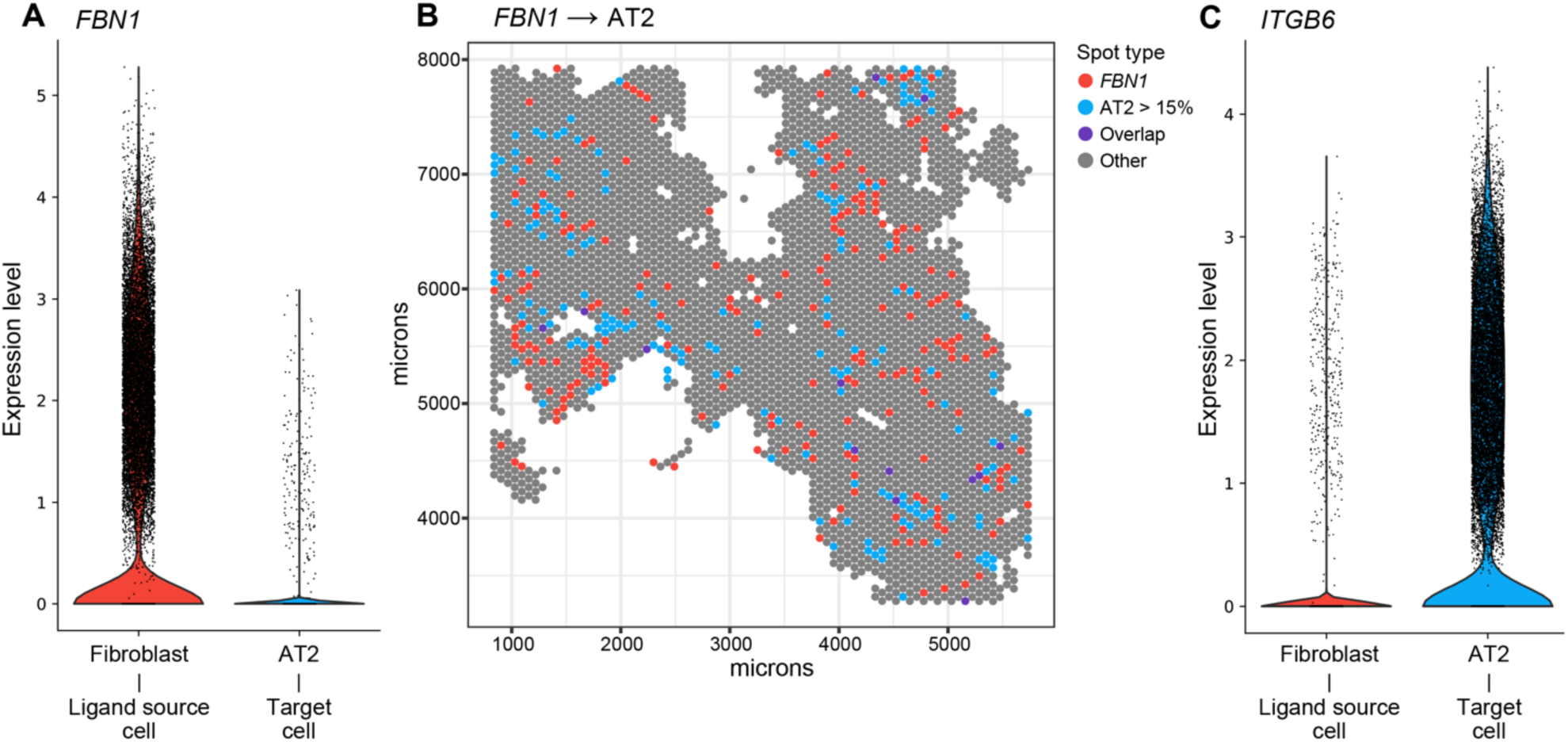
SPER detects non-overlapping paracrine signals (*FBN1*) in human lung. (**A**) Violin plots show expression levels (y-axis) of *FBN1* in relevant human lung cell types (x-axis) from reference scRNA-seq data. (**B**) ST plots visualize the spatial dependence between transcript-target cell type pairs *FBN1*/AT2 cells. The color of spots (color legend) shows whether *FBN1* is detected, a proportion of AT2 cell larger than 15%, both (overlap), or otherwise. (**C**) The expression levels of the relevant cognate receptor *ITGB6* in the reference scRNA-seq data.

## DISCUSSION

Sequencing technologies and computational methods for spatial transcriptomics are both developing rapidly, promising an exciting new wave of biological discoveries informed by spatial insight. However, despite the importance of cell type compositional changes, which are frequently the first analytical question asked in spatial and single-cell studies, and the potential of ST data to map them, and the well-established role of paracrine signals in dictating tissue composition [1], to our knowledge there has been no method to detect putative compositional regulators; SPER provides the first such method.

Although we performed evaluations using ST data generated with 10X Genomics Visium, SPER is not specific to this modality, and naturally generalizes to other spot-level technologies, such as Slide-seq [10] and XYZeq [45]. In these cases, the SPER workflow remains unchanged, requiring input ST expression data in the form of a genes-by-location matrix, spot locations, and estimated cell-type spatial compositional data from a deconvolution algorithm. For newer spatial technologies, such as CosMx (NanoString), MERSCOPE (Vizgen), and Xenium In Situ (10X Genomics), the spot-level and deconvolved cell-type compositional data is replaced by the cell-level spatially resolved data. This shift requires additional preprocessing for the data, treating every single cell as a spot with only one cell type. While the workflow remains functional, the computational burden for the single-cell data, often involving millions of cells, could be significantly higher than for Visium data, which typically includes two to four thousand spots. Since workhorse ST methods such as 10X Visium do not have single-cell resolution, analysis of their data using SPER requires estimating cell type compositions from ST data using any deconvolution method: SPER then models the compositionality of the spatial data using co-kriging, which can reduce the impact of noisy estimates while preserving core spatial patterns [18, 46]. Methods with higher deconvolution accuracy will increase the performance of SPER to capture biologically meaningful signals. Hence, we expect the SPER algorithm to be more powerful in the future as improved ST technologies and superior deconvolution approaches emerge.

Algorithms for paracrine signal inference such as SPER face two related challenges given the complexity of receptor ligand interactions. First, if a receptor-ligand interaction has been previously described, and SPER identifies it as highly significant in a particular dataset, since it is based on transcriptional measurements, this is only an inference, rather than definitive proof that the interaction is physiologically relevant. In this context, SPER operates as a discovery tool, generating hypotheses regarding the paracrine interactions that could be active in the tissue, and follow-up experiments are then necessary to translate these potential interactions into definitive new biological mechanisms. As we’ve shown, even a cursory inspection of the results of analysis by SPER in a well-known tissue (mouse brain), identifies both known and novel potential interactions. We thus expect that SPER can provide the first step in an analytical pipeline to discover active intercellular circuits that underly tissue structure, and if applied to perturbations or disease, that underly tissue transformation. A second challenge is that not all paracrine interactions are currently annotated—for this reason, SPER does not ignore putative regulatory transcripts that are unannotated as ligands or receptors, but rather enables a user to examine *all* highly ranked transcript-cell type pairs. In this way SPER can potentially uncover previously unknown intercellular signals that are associated with changes in tissue composition, providing the ability to identify regulators even, for example, if their receptor is unknown or is not detected due to lack of sensitivity.

Taken together, our results demonstrate that the application of SPER to ST data has the potential to identify which paracrine signals are involved in the intercellular circuits controlling homeostatic tissue composition, stress responses, tissue inflammation, and perhaps most importantly, pathological tissue transformations associated with disease. Identification of these signals is a critical step towards deepening systemic understanding of these complex processes and pinpointing novel biomarkers and potential therapeutic targets.

## METHODS

### SPER: Spatial Paired Expression Ratio

In general, SPER algorithm requires the normalized spatial transcriptomics data *E*_*n*×*m*_(the expression level of genes divided by their global average across all spots), the spots distance matrix *M*_*n*×*n*_, and the cell-type compositional data *C*_*k*×*n*_as the input, where *n* is the number of spots, *m* as the number of genes and *k* as the number of cell types. As the first step, the distance matrix of spots *M* is separated into *l* adjacency matrices *A*_*i*_, each recording whether two cells lie within certain distance range *d*_*i*−1_ ∼ *d*_*i*_ for *i* ∈ {0, 1, …, *l*} with *d*_0_ = 0. As a result, the distance matrix *M* can be estimated as the summarization of *d*_*i*_*A*_*i*_, where the original distances are binned into different groups. The default bin of the distance discretization is the distance between two spots, and all distances which are larger than the given largest distance limits are trimmed to that value. These adjacency matrices *A*_*i*_ ∈ Boolean^*n*×*n*^are normalized by multiplying a diagonal matrix *F*_*i*,*n*×*n*_, where each element is the reciprocal of each row’s sum of *A*_*i*_. Then, they are multiplied with the spatial transcriptomics data *E*_*n*×*m*_and the cell-type compositional data *C*_*k*×*n*_ as:

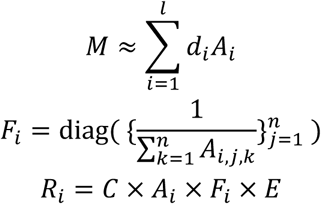

where *R*_*i*_ represents the *k* × *m* paired expression ratio matrix at the distance range *D*_*i*−1_ ∼ *D*_*i*_. Finally, a weight function **Φ** = (φ_1_, …, φ_*l*_) is applied on each gene/cell type pair across *R*_1_, …, *R*_*l*_. The default weight function follows a Poisson distribution Pois (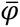) which has an extra penalty on overlapping and distal regions, where the parameter 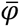 is chosen based on the potential functional distance of paracrine signal. Eventually, we can get the SPER score matrix *S* as:

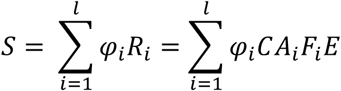

### Spatial ST data simulation

The simulations are created by the following steps: 1) A 80×80 grids is created and its distance matrix is calculated; 2) Cell type labels are assigned to each tile of the grid, which follows different mechanisms in three scenarios; 3) For each cell type, we create the corresponding marker genes and simulated their expression levels across all cells. Corresponding cell types will have higher counts of their markers with their counts following Poisson distributions Pois(*λ*) where *λ*∼Unif(15, 30). For other genes, including the non-marker and the control genes, their counts follow Poisson distributions Pois(*λ*′) where *λ*′∼Unif(0, 3). 4) The compositional data for each cell type is computed from marker genes’ expression level. The count of each cell type within a grid is sampled from another Poisson distribution. The final cell-type proportions are the fractions of each cell types within the tiles. Mathematically it can be shown as:

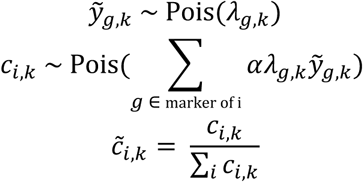

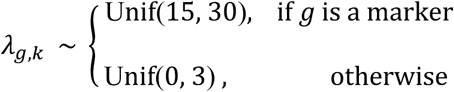

where ỹ_*g,k*_ is the expression level of gene *g* at tile *k*, 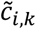 is the cell-type proportion of cell type *i* at tile *k*, and *α* is the scale factor.

Simplified single-source scenario simulates the source and target cells in a spatial relationship as ring (**Fig. 2**). Within the 80×80 grid, five cell types, two pairs of source and target cells and a control, are simulated. For each simulated cell type, we created 20 marker genes from two-dimensional normal distributions and calculated the cell-type spatial composition as described above. While the target cells have their marker genes simulated with given means and variances, the markers for source genes have their maximal values 10-unit away from the centers of target cell types. This spatial dependence characterizes the paracrine patterns between the source genes and target cells.

Realistic scenarios, both single-source and multi-source, we use Markov random field algorithm from the R package ‘mrf2d’ to simulate the spatial relationships between cells. Neighbors with L1 distance less than 3 are considered when calculating the 2D Markov random field. Source and target cells are set to spatially dependent on each other, while the target cells have a greater spatial autocorrelation. The single-source scenario simulated one pair of source and target cells and two control cell types, while the multi-source having two sources, one target, and two control cell types. The Pearson and Spearman correlation, and SPER scores are respectively calculated on three simulations.

We repeated each scenario 10 times given the random seed from 1 to 10. The resulting matrices from all repeats are used to predict the spatial dependence between genes and cell types, particularly the target cells and source genes. For each type, the ROC curve is plotted in distinguishing its designed paracrine signals from the other genes.

### Collection and pre-processing of scRNA-seq reference data

The scRNA-seq data were collected and pre-processed by Tasic, B., et al. in the Allen Institute for Brain Science, generated by the SMART-seq2 protocol [33]. The dataset contains 14249 adult mouse brain cortical cells and 34617 genes. The raw data was normalized by Seurat using SCTransform function [47], where 3000 subsampling cells were used to build the NB regression model and other parameters set as the default. The label ‘subclass’ was used as the annotation of cell type, which included 23 different subclass of mouse brain cells. Cajal–Retzius (CR) cell type, the count of which was only 7 in the scRNA-seq data, was removed from the data as the low cell counts weakened its statistical power. After this removal, 14242 cells with 22 distinct cell types were kept as the reference in the downstream analysis.

### Identifying cell-type-specific marker genes in scRNA-seq data

The cell-type-specific marker genes were found in the normalized scRNA-seq data by Seurat. For each cell type, their marker genes were then identified by Seurat using MAST one-versus-all test [34]. In these tests, each gene was tested whether it was differentially expressed in each cell type compared to rest of cells. Genes are selected as marker genes by the following criteria: fold change > 2, adjusted p-value < 10^-30^, fraction of expressing cells > 70%. All the other parameters were set as the default in the ‘FindMarkers’ function in the Seurat package [47].

### Collection and pre-processing of Visium spatial transcriptomics data in mouse brain

The mouse brain spatial transcriptomics data is collected from the 10X Genomics Visium public database [9]. It contains 2696 spots with 124367 mean reads per spot. The raw count matrix is normalized by Seurat using SCTransform function with default settings. The physical distances of cells are calculated from the provided spatial coordinates and the scale factor in the metadata. In the downstream analysis where ST and scRNA-seq data are needed, we only kept the genes detected in both datasets and having the total UMI counts greater than 40 in the normalized ST data.

### Cell-type deconvolution and co-kriging modeling in Visium mouse brain data

In the Visium mouse brain example, we used the normalized ST data to perform the cell-type deconvolution. The cell-type spatial compositional data *C* is estimated using RCTD full mode [11] with a reference using the scRNA-seq reference data. We select top 10 cell types which have the highest prevalence among all spots in the compositional data generated by RCTD. They are L2/3 IT, L5 IT, Meis2, L4, L6 IT, Oligo, Pvalb, L6 CT, Sst, and L6b. The other cell types are summed together as ‘Other’ aside from those 10 types. The cell-type spatial compositional matrix is further denoised using the co-kriging algorithms. 15 distance bins, 120 μm each, are used to compute the log-ratio variograms. To model the variograms between cell types, we use a compositional linear model of spatial coregionalization that includes a nugget effect term and 3 Gaussian variogram terms whose respective effective as 600, 1300, and 3000 micrometers. These 3 terms respectively represent the short-range, middle-range, and long-range interactions between cell types.

### SPER calculation

In the Visium mouse brain and human lung example, we used the normalized ST data and cell-type spatial compositional data as the SPER inputs. Calculation was performed using SPER R package. The weight function is derived from a Poisson distribution (λ=2.5) to give penalty on overlapping and distal regions while keeping its sum as 1, where the Poisson parameter 2.5 represents 180 nm in reality as the assumed average functional distance of paracrine signals. The choice of weight function is not unique, as users can manually fine-tune it to match the desired functional distance of the signals.

The final SPER score 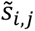 was calculated as:

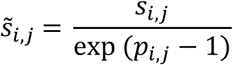

Where *S*_*i,j*_ stands for the raw SPER score and *p*_*i,j*_ for the expression prevalence of gene *i* and cell type *j*.

### Collection of extracellular gene set and ligand-receptor pair data

To identify the subcellular locations, we used the all channels integrated version of mouse extracellular gene set that includes knowledge, experiments, text mining, and predictions channel from the COMPARTMENTS database [48]. To filter out the intracellular genes, we select the genes whose locations are in the extracellular region (including extracellular space, matrix, exosome, and vesicle). Also, we only kept the genes whose confident level greater than 3. As a result, 4797 piece of genes’ information and 2622 different genes are left as the set of extracellular genes. After doing the intersection with the ST data, 1784 genes are included in the final extracellular gene set. The ligand-receptor interaction dataset is collected from FANTOM5 ligand-receptor datasets [49] and CellPhoneDB database [50]. Since all these databases only have the human genome data, we transformed the gene ID into the homologous gene in mouse genome to compare with the ST data we have got. There are totally 1888 ligand-receptor pairs that have the homologous genes in mouse, where 1636 of them are from FANTOM5 and 252 of them from CellPhoneDB.

### Calculating correlations and EMD of gene-cell type pairs

The Pearson and Spearman correlations are calculated between the spatial compositional data after co-kriging modeled and the spatial transcriptomics data. The same matrices, but size reduced, are used to calculate the EMD in comparing the computing complexity. We use the physical distance between spots as the ground distances, and the proportions of each cell-type composition are inputted as the weights when comparing to a gene’s spatial distribution. R package ‘emdist’ is used to calculate EMD in the simulations.

### Spatial-dependent signal detection

The SPER score matrix, Pearson correlation matrix and Spearman correlation matrix are used to filter the candidate genes and spatial-dependent signals with the help of scRNA-seq data as the reference of expression levels. When studying the enrichment of extracellular and ligand genes, we used the hypergeometric test to see if they are enriched within the gene set given thresholds. In both cases, the population size is 13436; the number of successes in the population is 1784 for the extracellular and 1888 for ligands. The sample size and the number of successes in the sample are dependent on the expression-level and measurement-level thresholds. If the p-values calculated from the tests are less than 0.05, we will take it as significantly enriched.

### Ablation study of co-kriging

We conducted an ablation sensitivity analysis to explore the contribution of co-kriging in modeling the cell type compositional data and SPER’s sensitivity. Following the same procedure as previously (**Fig. 4**), we applied SPER directly on the raw compositional data estimates generated from RCTD, with and without co-kriging to stabilize and model the compositional estimates. We used both statistical enrichment of extracellular cell location and participation as a known paracrine ligand as metrics to benchmark the performance with and without co-kriging (**Fig. S4**). We reasoned that after co-kriging modeling, SPER would be more likely to detect transcripts meeting these criteria, because of the ability of co-kriging to remove noise in compositional estimates. Consistent with this hypothesis, we observed that co-kriging led to an increase in the number of detected transcripts across most cell types, with out of 10 cell types exhibiting a higher number of transcripts with extracellular localization and 8 out of 10 showing an increase in putative paracrine ligands.

## Acknowledgements

The authors thank Leslie Gaffney for preparing and generation of figures for this article, Evan Lemire for programming assistance, and Yu Sun for helping improve the data visualization.

## Author contributions

T.Z. designed and implemented the SPER algorithm, performed data analysis and visualization and wrote the manuscript. A.H. conceived the study and wrote the manuscript.

## Declaration of interests

The authors declare no competing interests.

## Data and Code availability

An R package to implement the SPER algorithm is available for download at github.com/TianxiaoNYU/SPER. Extracellular gene set in mouse is available from the COMPARTMENTS database: https://compartments.jensenlab.org/Downloads; scRNA-seq reference data is obtained from Allen Institute for Brain Science: https://portal.brain-map.org/atlases-and-data/rnaseq; Visium spatial transcriptomics data from mouse brain is available from 10X Genomics dataset: https://www.10xgenomics.com/resources/datasets/mouse-brain-serial-section-1-sagittal-anterior-1-standard-1-0-0. An R markdown tutorial to help generate the main results of this article using SPER package, as well as all other necessary data files in this study can be found at: https://www.dropbox.com/scl/fo/pep07jjid71rr72voeeik/h?rlkey=2agyojo1ugf5l61bqtal6hjy0&dl=0.

## SUPPLEMENTARY FIGURES

**Supplementary Figure 1.**
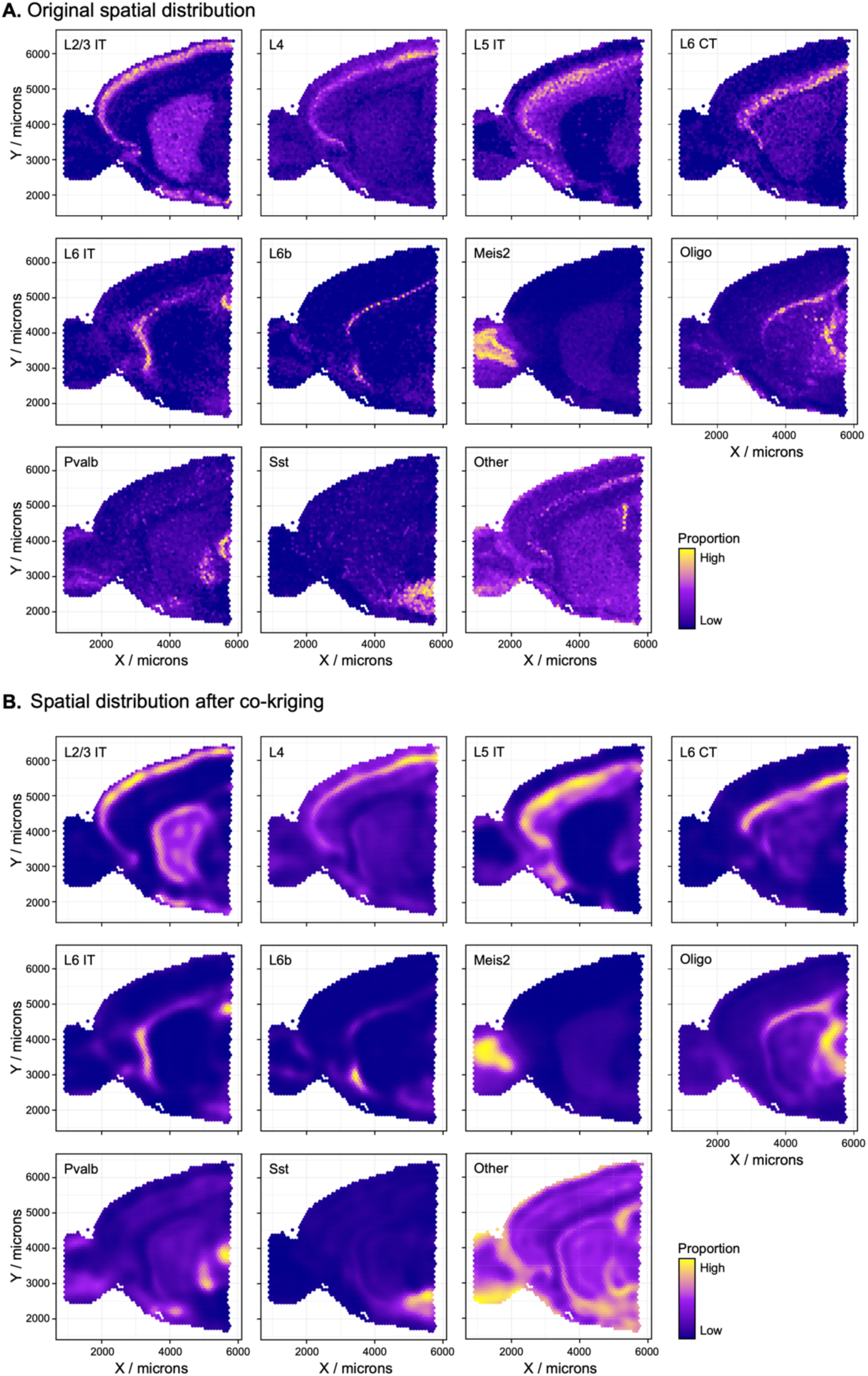
Co-kriging enables SPER to effectively denoise spatial compositional data. Spatial distribution of composition of primary cell types decomposed by RCTD in full mode before (A) and after co-kriging modeling (B). The top ten most abundant cell types are modeled individually, and the other 11 cell types are summed as the ‘Other’ category. In each subplot, X and Y-axis represent the physical coordinates of spots, and the color represents the normalized proportion of the relevant cell type.

**Supplementary Figure 2.**
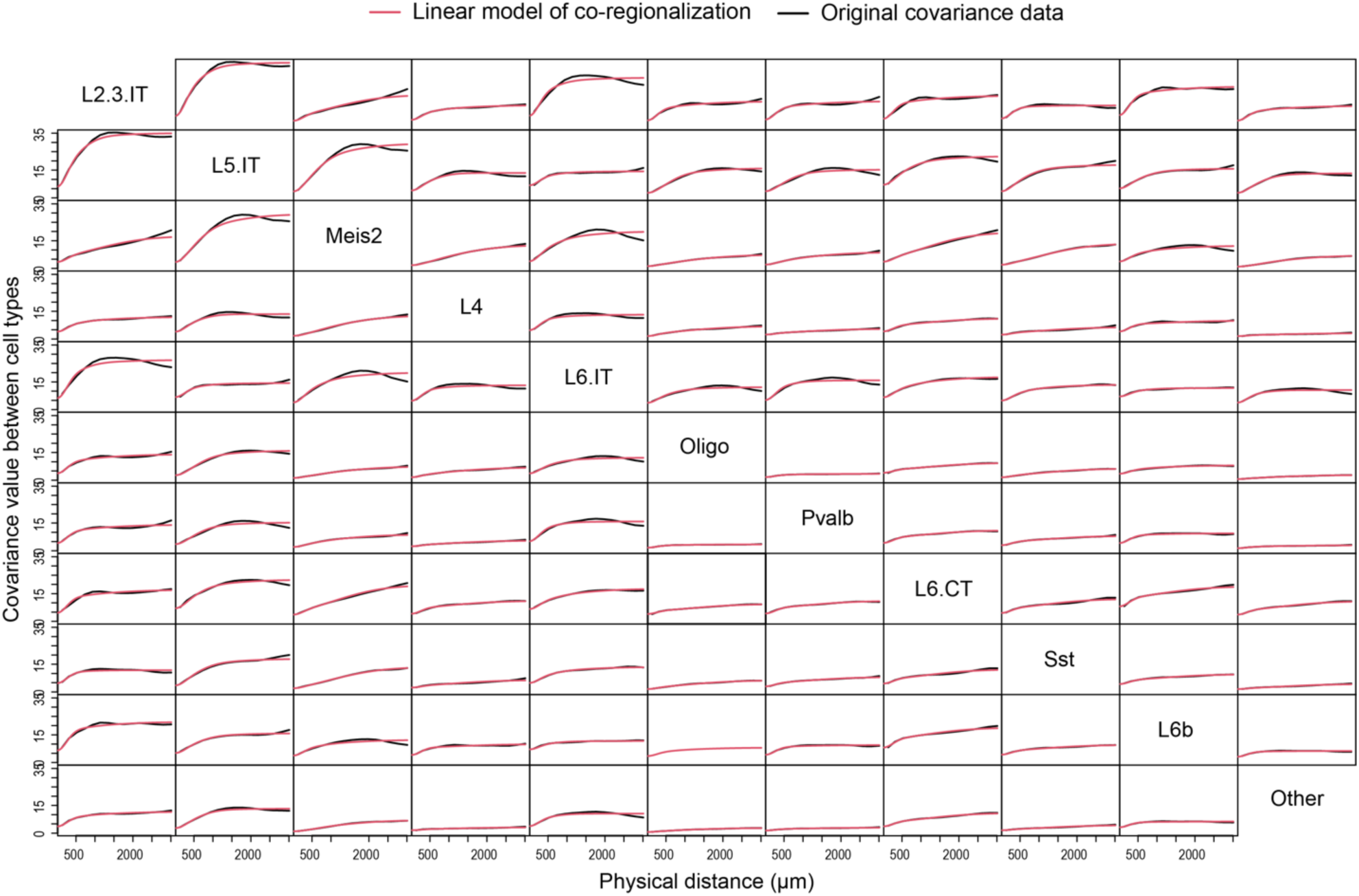
Co-kriging fits visualized as variograms. Co-kriging method fits on the ten neuronal cell types from an example dataset from mouse brain. A compositional linear model of co-regionalization (red line) with a ‘nugget’ effect term for short scale randomness and three Gaussian variogram terms to model effects at different ranges (600, 1300, and 3000 μm) showed excellent fit to the original covariance data (black line) collected and calculated at 15 uniformly distributed lags. The labels at the diagonal identify the cell type for each row and column, while ‘Other’ represents the sum of all remaining cell types. The x-axis in each subpanel represents the physical distance in μm, and y-axis represents the relative value of the covariance between each pair of cell types.

**Supplementary Figure 3.**
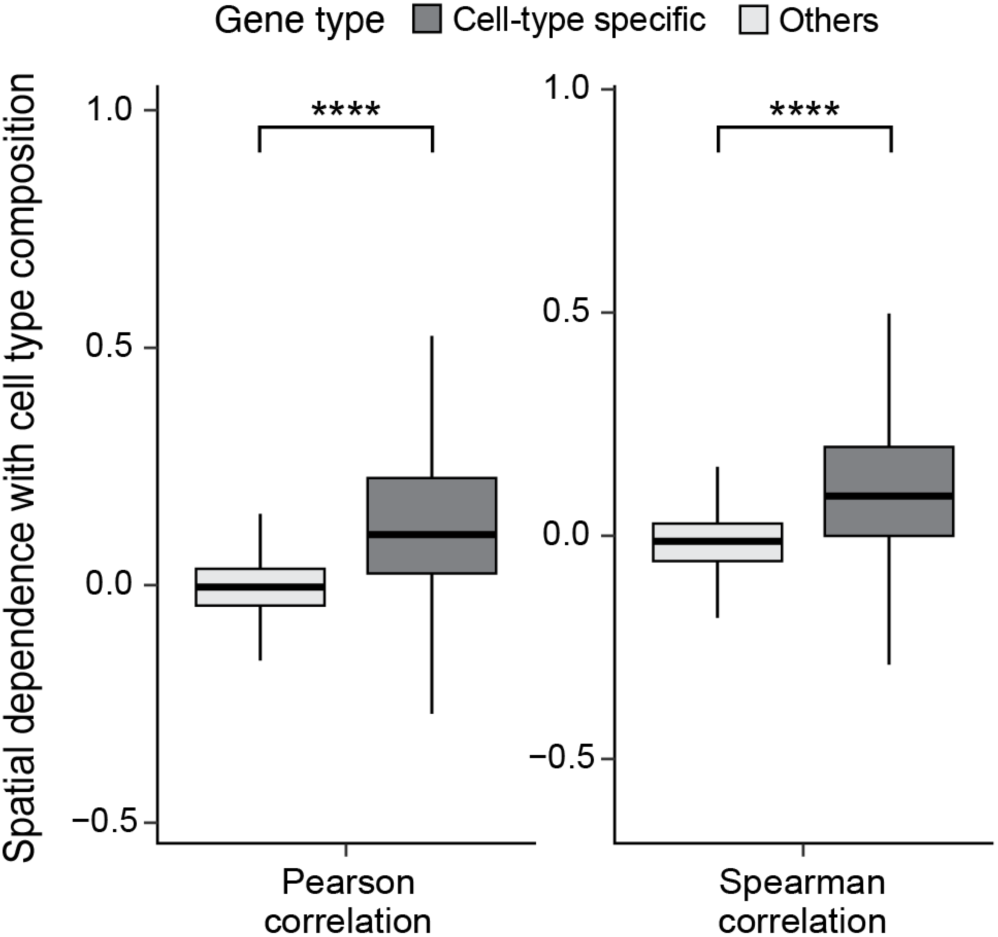
Pearson and Spearman correlations detect marker genes with overlapping spatial dependency. Boxplots showing Pearson and Spearman correlations between cell type spatial compositions and gene’s spatial distribution. Marker genes for each cell types are selected from the scRNA-seq reference data by MAST. The four-star labels represent the difference (adjusted p-value < 10^-4^) in the measurement scores between the marker and non-marker groups.

**Supplementary Figure 4.**
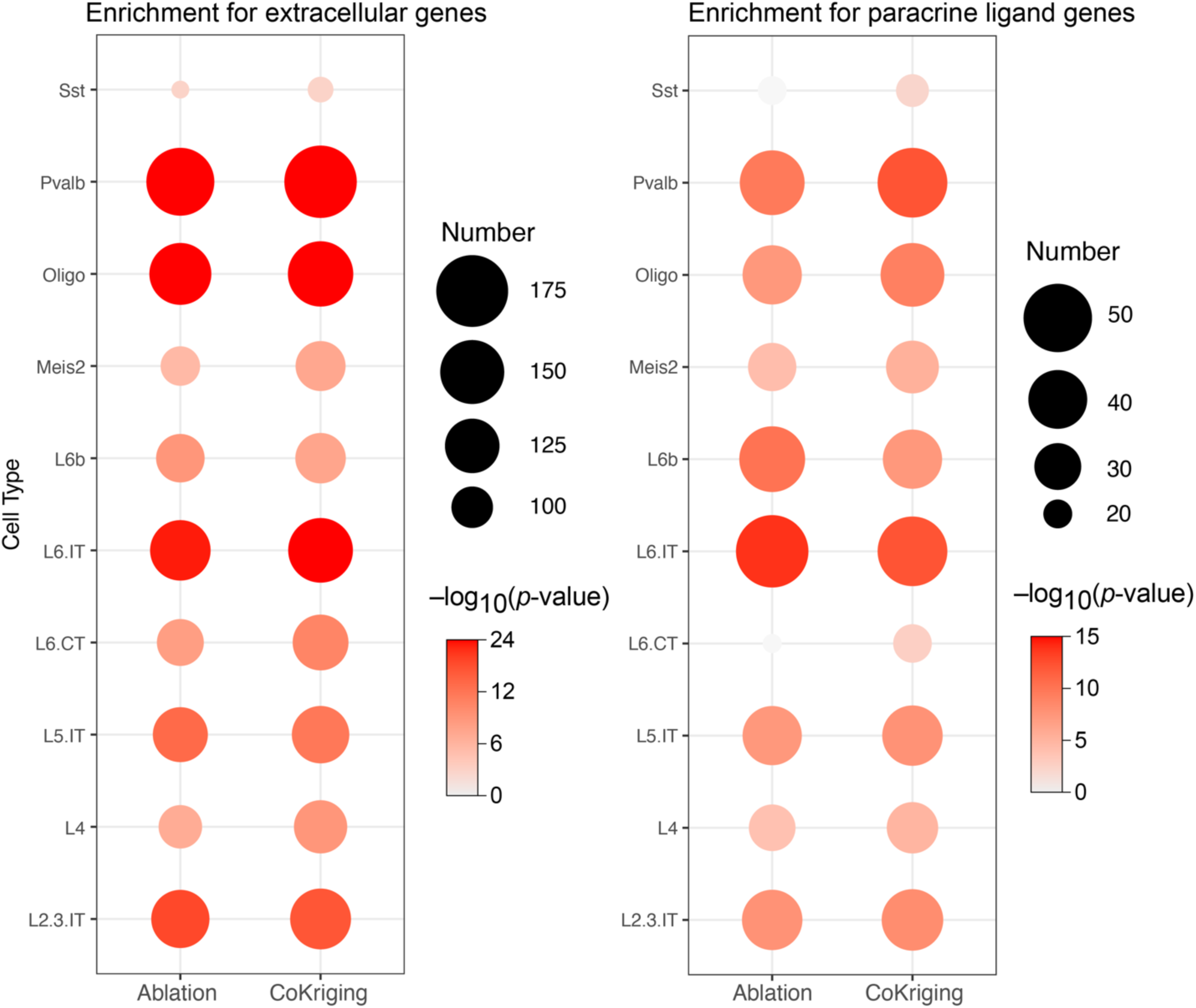
Ablation study demonstrates SPER detect more paracrine ligand genes with co-kriging. Dot plots visualize hypergeometric enrichment analysis of known paracrine ligand signals with and without co-kriging analysis. Ligand genes annotations were collected from COMPARTMENTS, FANTOM5, and CellPhoneDB databases respectively (see **Methods**). The candidate sets include genes whose scores for each spatial dependency metric (x-axis) are above than the 95% percentile of all genes. The size of dots represents the number of extracellular or ligand genes in the given gene set. The color represents the significance the enrichment in the set (hypergeometric test), non-significant (*p*>0.05) overlaps are shown as light grey. With co-kriging, the number of transcripts in both categories increased for most cell types, with 9 out of 10 showing an increase for extracellular location and 8 out of 10 for paracrine ligands.

**Supplementary Figure 5.**
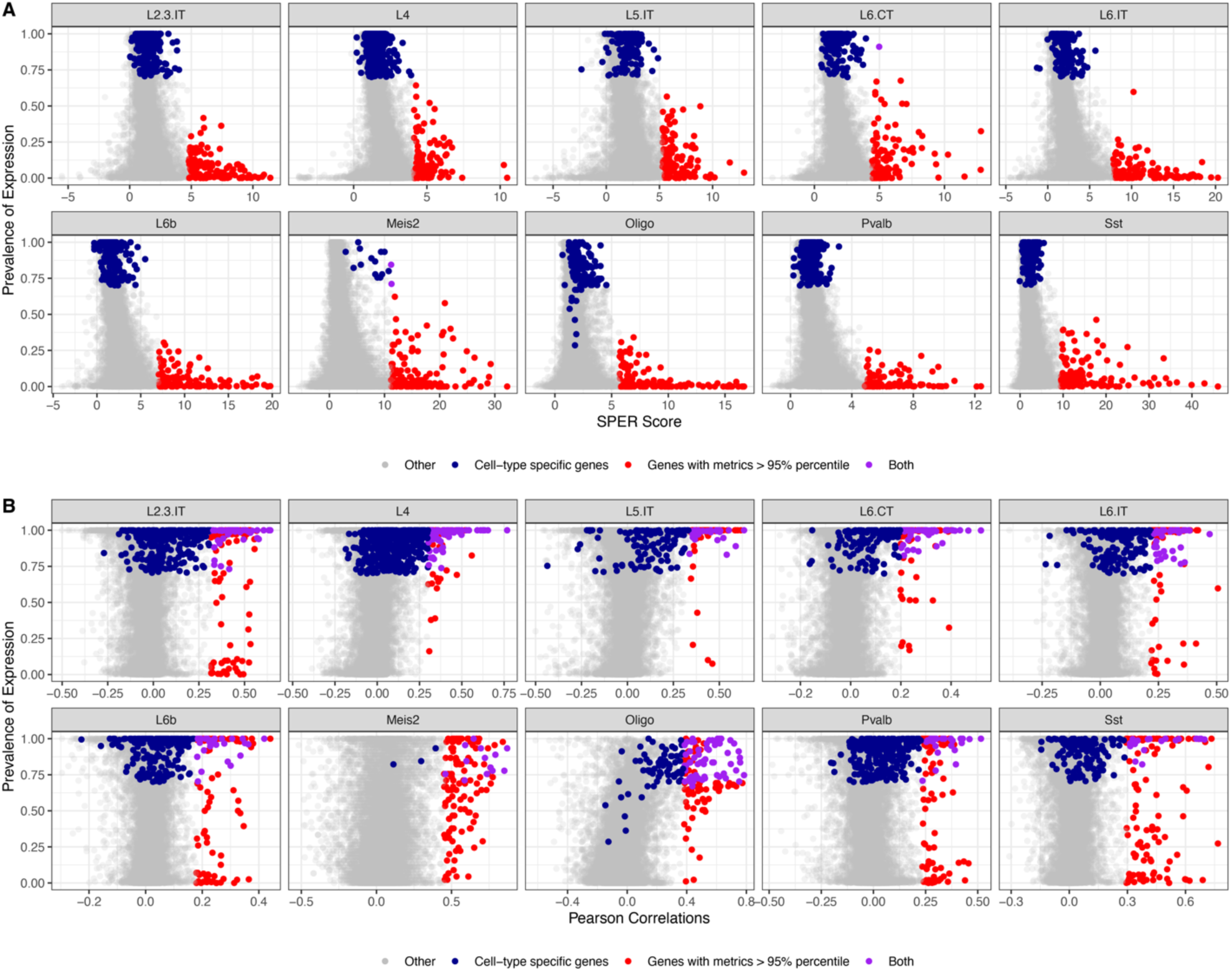
Comparison of trivial spatial associations and genuine putative regulators between SPER and Pearson correlation. Scatter plots show the distribution of SPER (A) and Pearson correlation (B) scores (x axis) with the proportion of cells of each type (plots) in which each gene (points) is detected (y axis) in scRNA-seq data. The dark blue dots mark the marker genes for each cell type (found by MAST, details in Methods). Red dots stand for the genes whose measurement scores larger than 95% percentile. Purple dots represent genes that meets both conditions.

## Notes

### Competing Interest Statement

The authors have declared no competing interest.

### Summary of Updates

Manuscript text and figures revised and updated, incorporating new data and results.

